# An intrinsic neural eye tracker in primary visual cortex

**DOI:** 10.1101/443507

**Authors:** Adam P. Morris, Bart Krekelberg

## Abstract

Humans and other primates rely on eye movements to explore visual scenes and to track moving objects. As a result, the image that is projected onto the retina – and propagated throughout the visual cortical hierarchy – is almost constantly changing and makes little sense without taking into account the momentary direction of gaze. How is this achieved in the visual system? Here we show that in primary visual cortex (V1), the earliest stage of cortical vision, neural representations carry an embedded “eye tracker” that signals the direction of gaze associated with each image. Using chronically implanted multi-electrode arrays, we recorded the activity of neurons in V1 during tasks requiring fast (exploratory) and slow (pursuit) eye movements. Neurons were stimulated with flickering, full-field luminance noise at all times. As in previous studies ^1-4^, we observed neurons that were sensitive to gaze direction during fixation, despite comparable stimulation of their receptive fields. We trained a decoder to translate neural activity into metric estimates of (stationary) gaze direction. This decoded signal not only tracked the eye accurately during fixation, but also during fast and slow eye movements, even though the decoder had not been exposed to data from these behavioural states. Moreover, this signal lagged the real eye by approximately the time it took for new visual information to travel from the retina to cortex. Using simulations, we show that this V1 eye position signal could be used to take into account the sensory consequences of eye movements and map the fleeting positions of objects on the retina onto their stable position in the world.

---

During everyday vision, fast eye movements known as ‘saccades’ shift the line of sight to place the high-resolution fovea onto objects of interest. These movements are complemented by slower, smooth eye rotations used to track moving objects (e.g. a bird in flight) or to lock gaze on an object during locomotion^5^. Though essential, these movements have stark consequences for the retina’s view of the world: objects that are stationary appear to jump or move, and objects that are in motion travel along grossly distorted trajectories (Fig 1). These phenomena reflect the fact that visual input arrives to the nervous system in an *eye-centred* (or retinal) coordinate system, in which objects evoke activity within a local patch of photoreceptors and ganglion cells (i.e. as a place code). Our perception of the world, however, is entirely different: objects do not appear to change positions every time we move our eyes, and we have no trouble planning actions toward them. This suggests that the visual system uses internal knowledge of eye position (and head position in the case of combined eye-head gaze shifts) to compute the *true* locations of objects from their fleeting retinal projections (i.e. their positions in head- and/or body-centred coordinates). Where, and how, this transformation take place in the brain remains unclear.

**Figure 1.**
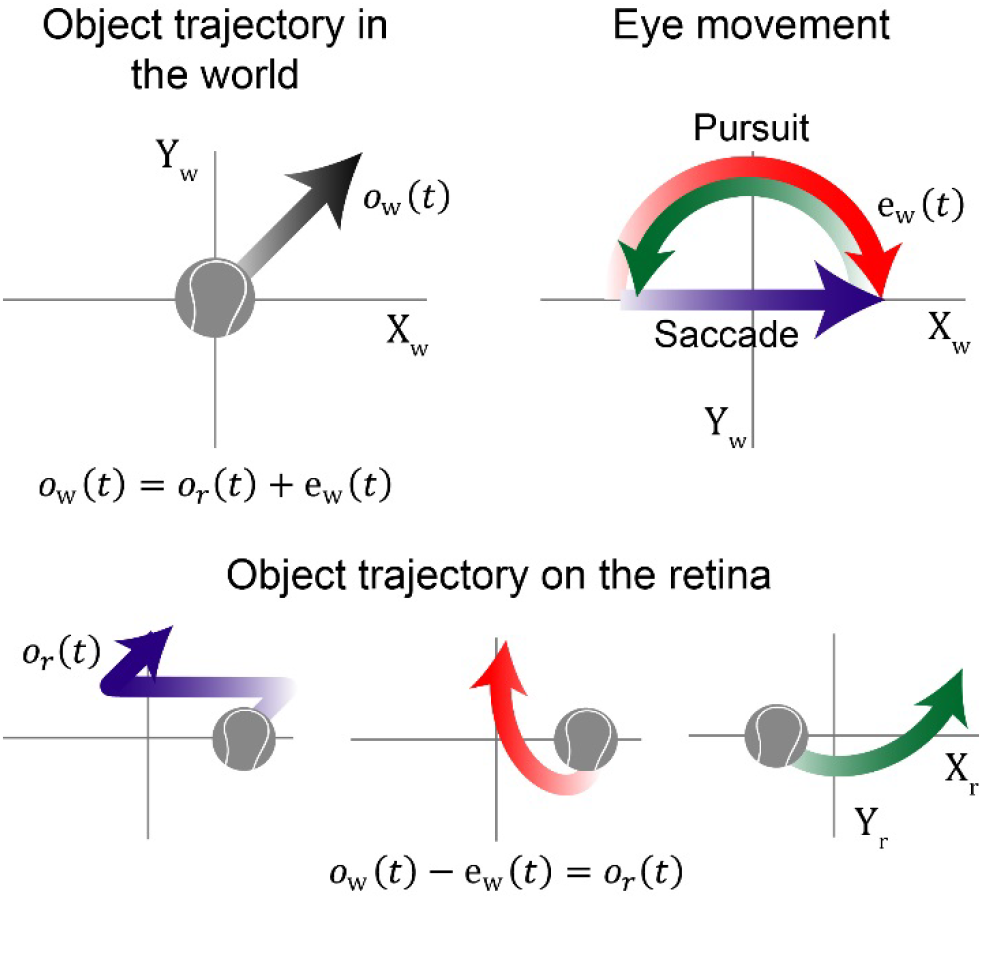
Our sense of a stable visual world arises from retinal images that change with every eye movement. In the cartoon, a fixed state of the world (i.e., a ball in motion at 5°/s; black arrow) generates three different patterns of retinal stimulation (lower panels) under different eye movement scenarios: a fixate-saccade-fixate sequence (purple arrow), and smooth pursuit at 10°/s in opposite directions around a half-circle (green and red arrows). Note that different coordinate systems are used across panels: world-coordinates (*X_w_*, *Y_w_*) for the true object and eye, and retinal coordinates (*X_r_*, *Y_r_*) for the visual images (hereafter, these *r* and *w* subscripts are used to indicate the coordinate system of a spatial variable). The object’s trajectory on the retina, *o_r_*(*t*), is roughly equal to the object’s trajectory in the world, *o_w_*(*t*), minus the path travelled by the eyes, *e_w_*(*t*). The brain, however, must make the reverse inference to recover the object’s true trajectory, i.e., *o_w_*(*t*) = *o_r_*(*t*) + *e_w_*(*t*). Here, we show that V1 neurons represent in parallel both terms on the right side of this equation.

At first glance, it would seem that a stable representation of visual space is not achieved until late in the cortical hierarchy. For example, in the first stage of cortical vision – the primary visual cortex (V1) – neurons have receptive fields (RFs) that are fixed on the retina ^6^, and so move across a visual scene with the eye. Patterns of activity within the V1 retinotopic map thus reflect what is happening on the retina, not in the world itself. Perhaps surprisingly, this remains true throughout the cortex beyond V1, where many visually sensitive neurons have RFs that remain anchored to the retina ^7-9^. Similarly, neurons that are sensitive to image motion tend to prefer the same direction on the retina regardless of whether it was caused by a moving object or a moving eye ^10, 11^.

Other neurons within the parietal and temporal lobes, however, do show a greater degree of stability, such that their selectivity shifts (in retinal coordinates) with eye movements to compensate for the retinal displacement ^11-16^. Their selectivity is thus at least partially anchored in head- or world-coordinates. These observations, combined with other known forms of spatial remapping during eye movements ^14, 17^, support the idea that stability is achieved only late in the visual processing stream; and even then, only within the dorsal “where” pathway that supports the planning of actions (see ^18^ for a review). In this view, the ventral “what” pathway that supports recognition of faces and objects must rely on its dorsal stream counterpart to provide the spatial information necessary to piece together snapshots taken from different lines of sight.

We tested the alternative hypothesis that visual stability begins in V1 – despite superficial appearances to the contrary – through a distributed population code ^19, 20^. Such a solution would have the advantage that all downstream computations – in both the dorsal and ventral streams – have direct access to visual information that is robust to eye movements. Our hypothesis builds on previous reports of neurons throughout visual cortex, including in V1, that alter their rates of action potentials (spikes) systematically with changes in eye position, even when the image within their RFs is held constant – a property known as a ‘gain-field’ ^1-4, 21, 22^. For such neurons, changes in eye position do not alter the selectivity to the retinal image, but do influence the *gain* of the visual response (e.g. the amplitude of the tuning curve) and therefore carry information about gaze direction. This is critical, because to a first approximation, the location of an object in a scene is simply the sum of its position on the retina and the current eye position (we ignore head movements and other postural changes; Fig 1). Thus, a population of gain-field neurons might allow an eye-centred area like V1 to nevertheless carry a stable representation of visual space. This prediction is supported by influential computational studies ^19, 20, 23^ that incorporated artificial, idealized gain-field neurons, but has to our knowledge never been tested empirically.

For this hypothesis to be true, real gain-fields in V1 must support an accurate representation of the position of the eye during natural vision. We and others have shown that neurons in *higher* areas of the dorsal pathway reliably code the direction of gaze during fixation ^24-26^, and predict the landing point of an impending saccade ^25, 27, 28^. Here we show that not only is a reliable eye-position signal present much earlier – in V1 – but also that it can be incorporated into downstream computations in the same way irrespective of whether the eyes are exploring a scene (i.e., saccades) or tracking an object (i.e., pursuit). Thus, this crucial ingredient for visual stability is available to both the dorsal and ventral processing streams for perception and action.

## Results

We recorded extracellular action potentials from 397 units (232 single-units and 165 multi-units) using multi-electrode arrays permanently implanted in the parafoveal region of V1 in two adult macaques (Fig 2). There were no qualitative differences between single and multi-unit responses (see below) hence we analysed them together. On ‘saccade’ trials (included in all recording sessions), the animal fixated a target at one of eight possible starting positions arranged in a circle (15° diameter; only one target was displayed at a time). After a delay of 1100 ms, the target stepped to the opposite side of the circle, requiring an immediate saccade and fixation at the new position for a further 1100 ms. ‘Pursuit’ trials (interleaved with saccade trials in 45/68 sessions) were identical except that after 500 ms of initial fixation, the target moved clockwise (CW) or counter clockwise (CCW) around the imaginary circle to the opposite side (speed = 10°/s), requiring the animal to track its position with pursuit eye movements. A final trial type (all sessions) required fixation on a central target for the whole trial.

**Figure 2:**
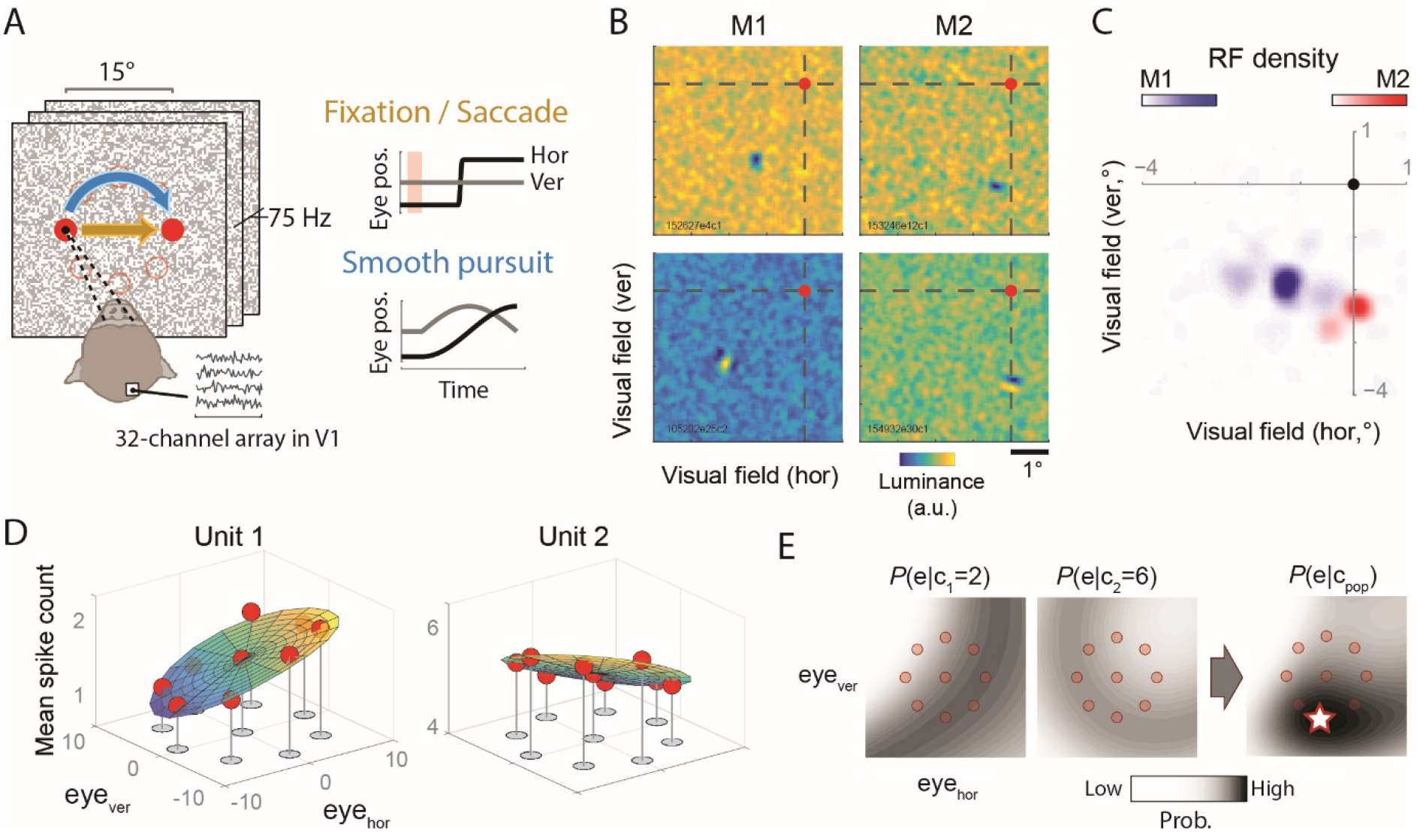
(A) Experimental design, including idealized eye traces for saccade and pursuit tasks (9 o’clock starting position shown). The open circles show the other possible starting positions for the task. The pink shaded region (−700 to −300 ms relative to saccade onset) was defined as the fixation epoch. (B) Receptive field estimates (i.e., spike-triggered averages) for two example V1 units (rows) from each animal (M1 and M2), plotted relative to the fovea (red dot). (C) The distribution of receptive fields across all recordings. (D) Example gain-fields for two units, plotted as mean spike counts for 100 ms samples during the fixation epoch at each of nine eye positions (the standard errors are omitted for clarity, but were smaller than the symbols in all cases). Under the assumption of Poisson variability, the fitted regression surfaces provide a generative model relating eye position to spike-count probabilities – a ‘probabilistic gain-field’ (Methods). (E) Population decoding using probabilistic gain-fields. The two leftmost images show the probability of all possible eye positions (across a finely spaced 2D grid) given observed spike counts of 2 and 6 for units 1 and 2 in (D), respectively. The rightmost image shows the combined probability map for this minimalistic population of two units. The position of the peak (star) represents the maximum likelihood estimate of eye position for this example, [ê_wx_. ê_wy_]. In practise, the decoded population included all units with gain-fields (see Methods).

Throughout each trial, neurons were stimulated visually by flickering noise in the display background, achieved by setting every pixel to black or white randomly every 13.3ms. The high spatiotemporal frequency of the noise ensured comparable stimulation of RFs regardless of eye position or velocity (except during the saccade itself, where high velocity retinal slip would have caused grey-out). Further, it allowed the use of spike-triggered reverse-correlation to estimate the RFs of the recorded units. Using this approach, we confirmed that the observed RFs were located within the lower parafoveal region in both animals (1.5-3° from the fovea; Fig 2B, C).

### Decoding eye position during fixation

We first extracted spike counts during the initial fixation epoch on saccade trials (Fig 2A; 100 ms windows), when the eyes were stationary at eight different positions, and for a matched epoch on the central fixation trials. Consistent with previous reports ^1-3^, roughly half of the units (219/397; 127 single-units and 92 multi-units) showed significant differences in mean firing rate across the nine positions, even though the RF was stimulated by statistically identical visual noise – that is, they showed gain-fields (Fig 2D; significance was determined using stepwise regression; see Methods). The mean strength of this modulation was 43% (SD = 42%; median = 28%), defined as the difference between the maximum and minimum mean rate divided by the global mean. In control experiments, we confirmed that these modulations were not driven by differences elsewhere in the retinal image that co-varied with eye-position, in particular, the edges of the display (which were at least 6° outside of the RFs at all times; Supplementary Figure 1).

The presence of gain-fields hints that as a population, these units might carry enough information to encode 2D eye position (i.e. horizontal and vertical orientation of the eye in the orbit, or equivalently, fixation position on the screen). We developed a decoder based on maximum-likelihood estimation (Fig 2E). The traditional definition of a gain-field quantifies only how the mean firing rate of a neuron depends on eye position and says nothing about spike-count variability. We introduced a richer statistical quantification known as a *probabilistic gain-field* (pGF) ^24^, which quantifies the likelihood of observing each possible spike-count (i.e., *c* = 0, 1, 2, 3, etc.) at all possible eye-positions (i.e. the pGF is a generative model for spike-counts during fixation, even for positions not actually tested in the experiment). Using Bayes rule, and the assumption that all eye positions are equally likely *a-priori*, a collection of pGFs (one for each neuron) can be used to translate an observed population response (i.e. vector of spike-counts) into an estimate of the current eye position, [ê_wx_. ê_wy_].

Because the 219 units were not actually recorded simultaneously, the decoding analysis was based on a *pseudo*-population response approach. That is, we sampled a trial at random for each neuron (from the pool of trials available for a given fixation position) and compiled the spike counts into a population response over time. We then decoded the pseudo-population response at each time point (independently). This was repeated for 1000 pseudo-trials for each condition (Methods).

Figure 3A shows the decoder performance for population responses measured during the fixation epoch (using cross-validation on a data-set not used to setup the decoder). V1 represented the eye accurately and precisely across all of the tested positions, albeit with slight spatial compression (mean *constant error* across conditions was 1.29° [STE = 0.16]; *variable error*, defined as the median Euclidean distance of the trial-by-trial estimates from the distribution median, and averaged over conditions was 3.35° [STE = 0.32°]). Comparable performance was observed when we estimated the initial fixation position on pursuit trials (Fig 3A), albeit with greater error (mean constant error = 1.97° [STE = 0.32°]) due to a smaller population size (pursuit trials were included during recordings for only 179/219 units). As a further test of robustness, we decoded the fixation epoch *after* the saccade (+200 to +500 ms) and again found good accuracy (mean constant error = 1.82° [STE = 0.23°]), even though the decoder had never been exposed to data from these time points.

**Figure 3:**
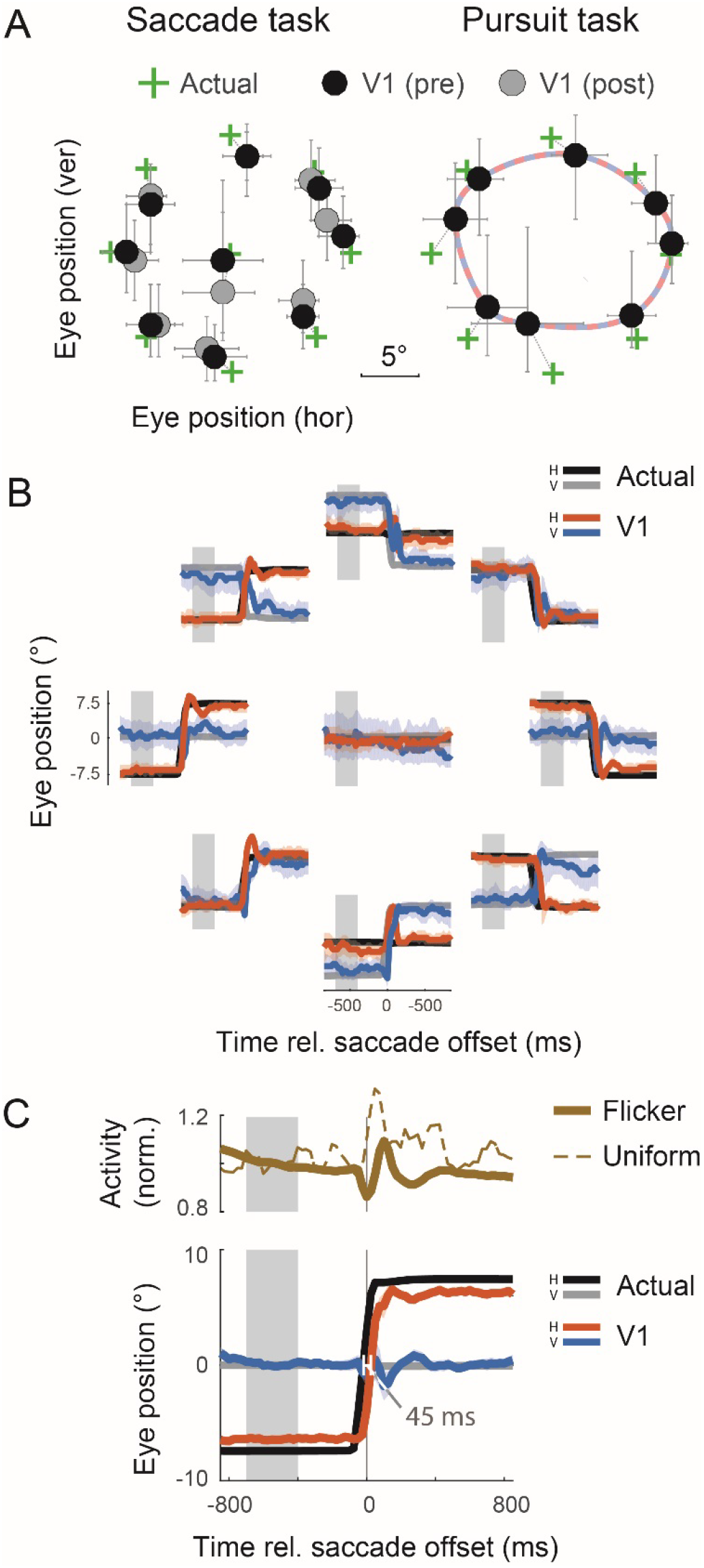
The representation of eye position in V1 is accurate during fixation and updates rapidly with each saccade. (A) Eye-position was decoded from 100 ms samples of neural activity. Left panel: median decoded eye positions (using the activity of 219 units) during fixation before (“pre”) and after (“post”) the saccade, compared with the actual fixation positions. Error bars represent variability (middle 50% of the distribution) across samples (trials and time). Right panel: decoded positions (using the activity of 179 units) during initial fixation on pursuit trials (i.e., −200 ms to +100 ms relative to target motion onset). The dashed line indicates a cubic spline 2D interpolation used to benchmark decoder performance during pursuit (see Fig 4). (B) The decoded eye position over time for each saccade direction (red and blue curves), plotted against the average eye trace (black and grey). Plots are arranged spatially according to the starting position (no saccade was required for the central fixation position). Curves represents the median decoded position and shaded errors represents the variability (middle 50% of distribution) across trials. The grey shaded region indicates the fixation epoch used to build the decoder (i.e. to estimate a pGF for each neuron). (C) The data from (A), averaged over all eight starting positions (after rotating the data to align their starting positions). Error shading = ±1 STE. The upper panel shows the mean firing rate across the population (normalised to the individual rates within the fixation epoch) in response to the flickering background and for a uniform grey background (the latter was performed in only a sub-set of experiments; N = 61 units).

Taken together, these results show that primary visual cortex contains a robust representation of the eye during fixation. This is similar to our previous application of this decoding approach to neural data recorded in extrastriate and parietal cortices ^24^.

### Decoding eye position during fast movements

Next, we asked whether this decoder, which was based only on the gain-fields measured during fixation (i.e., no re-training was performed), could also track the eyes across saccadic and pursuit eye movements. For this to be the case, firing rates must continue to be influenced by the eyes during these other oculomotor behaviours and remain statistically consistent with those observed during fixation. For example, if a neuron showed higher mean firing rates during fixation on the right side compared with the left, it must similarly do so in the immediate wake of a saccade to this position, and when the eyes pass smoothly through it during object tracking.

To reveal the representation of eye position during saccades in V1, we applied the fixation decoder sequentially and independently to each point in time throughout the fixation-saccade-fixation sequence (Fig 3B, C). As we have already seen, the decoder estimated accurately the initial and final eye positions. The transition between these points – the neural “saccade” – occurred shortly after the true change in eye position and with approximately sigmoid-shaped dynamics, much like the actual eye. This was true for all eight saccade directions and starting positions (Fig 3B). This consistency is reflected in the sharp dynamics of the decoded eye-position signal even when averaged over conditions (after first rotating them onto a common starting position; Fig 3C). Almost identical results were found when we repeated these analyses using only the single-units (127/219) in our sample (the correlation with the data shown in Fig 3C was 0.997; data not shown).

To estimate the onset and offset times of the neural saccade, and to compare them with those of the actual eye movement, we fitted the averaged eye-position signal and the eye-tracking data in Fig 3C with sigmoids (cumulative Gaussians; see Methods). Using parameters from the fits, we found that the V1 saccade lagged the actual eye by just 45 ms and had a similar duration (13 ms longer than the real eye). This short lag is well within the duration of a typical fixation ^29^, suggesting that the V1 representation of the eye is able to keep up with the eye during exploratory vision. A comparable signal delay was found using cross-correlation analysis (optimum lag = 46 ms; data not shown). At this lag, the decoded eye-position signal accounted for 97% of the variance in the true eye position over the course of the trial.

The stable, sigmoid-like dynamics of the V1 eye-position signal contrasts with the overall responsivity of the V1 population. The average firing rate showed complex modulations over time, including a strong reduction during the saccade followed by an enhancement shortly after, as well as more gradual declines over time during the fixation periods (the mean rate during the post-saccadic fixation was reduced relative to the pre-saccadic fixation by 6.0% on average [SD = 6.7%]; Fig 3C). The transient drop during saccades likely reflects in part “grey out” of the white-noise stimulus due to retinal slip and temporal integration. The post-saccadic enhancement, however, is unlikely to be a consequence of changes in the retinal image during the eye movement because it occurred similarly even for saccades across a uniform grey screen (dashed line), consistent with earlier reports ^30-33^. The timing of the enhancement coincided with the endpoint of the decoded V1 saccade, suggesting the arrival of a signal that underlies both enhancement and the updating of gain-fields.

The stability of the decoded V1 eye-position signal in the face of this variability in the global firing rate – to which the decoder had not been exposed previously – suggests that the population nevertheless retained appropriate patterns of activity across neurons. Invariances of this kind are among the key computational advantages of coding information using distributed population codes ^19, 20^.

### Decoding eye position during pursuit movements

The previous section showed that the transition from one fixation to another by a saccadic eye movement is represented faithfully in V1 (albeit with a short lag). Smooth pursuit, however, is a very different kind of oculomotor behaviour, both kinematically and in its neural control mechanisms ^34^. We tested whether the neural representation of the eye in V1 is nevertheless maintained during pursuit and can again be read out using our “fixation” decoder (because pursuit trials were included only in a subset of recording sessions, we decoded using only 179 of the 219 units).

In our task, the eye tracked the target along a circular path at a constant speed, passing through the positions measured during the fixation task on other trials (which we will refer to as ‘waypoints’). The horizontal (*e_wx_*) and vertical (*e_wy_*) coordinates of the eye therefore traced out a (half-period) sinusoid over time with an amplitude of approximately 7.5° (Fig 2A).

Figure 4 plots the decoded eye position during pursuit for each of the eight starting positions (CW only; Fig 4A) and averaged over conditions (separately for each pursuit direction; Fig 4B). The decoder tracked the position of the eye tightly in space and time on average, as it did for saccades, but with a centripetal bias toward the primary (central) gaze position. We quantified this performance with simultaneous sine and cosine fits to the horizontal and vertical components of the decoded eye position (not shown in the figure; see Methods). The amplitude of the fit showed that the decoded signal had a reduced pursuit radius of 5.62° (i.e. 76% of the true value).

**Figure 4:**
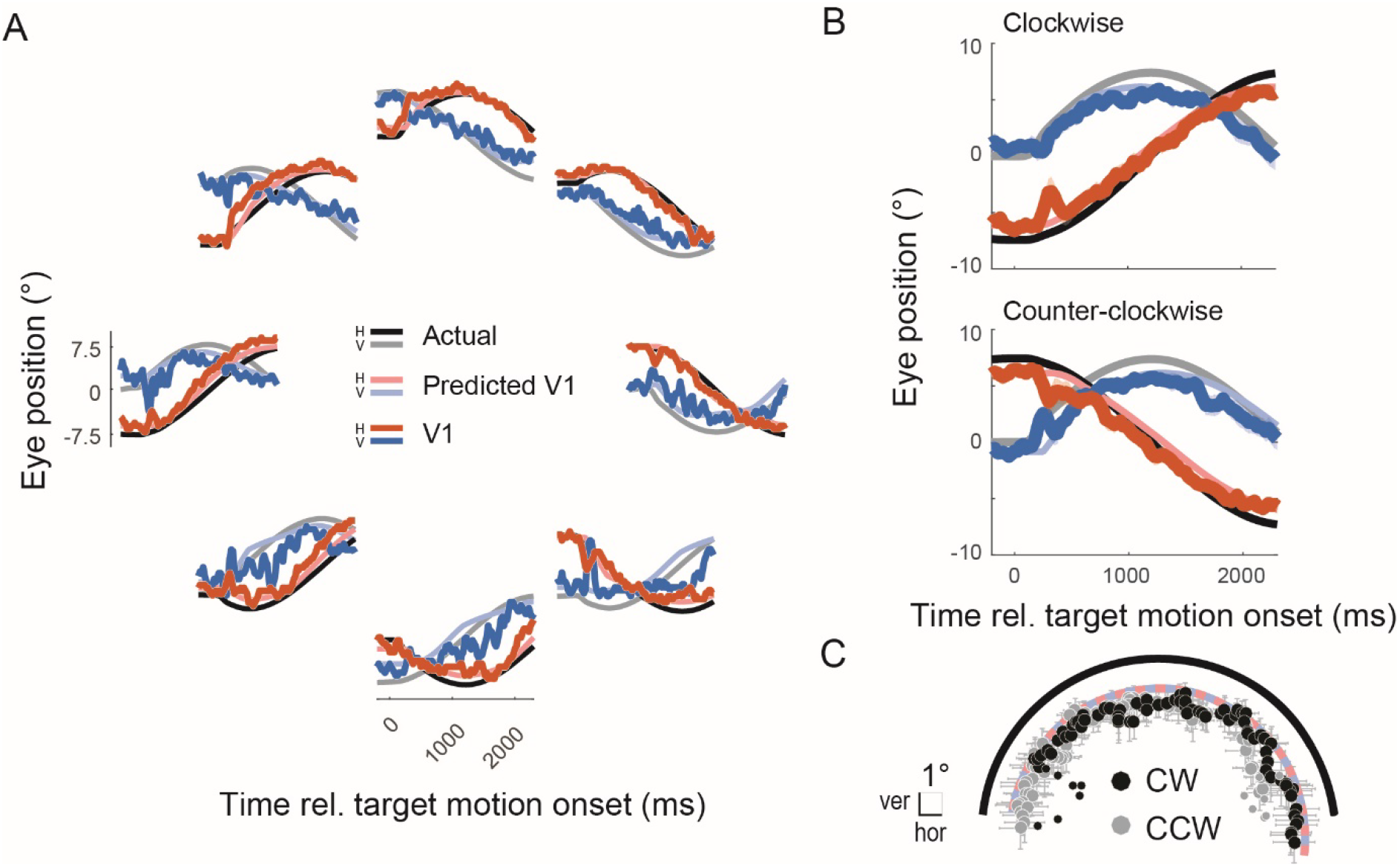
V1 neurons track the eye during pursuit using the same neural code as during fixation. (A) Decoded eye position over time for clockwise pursuit, plotted as in Figure 3B, but with the addition of curves showing what the decoder performance would look like if the spatial errors in decoder performance seen during fixation (Fig 3A) were recapitulated during pursuit (“predicted V1”). The predicted V1 signal shown here corresponds to the cubic spline shown in Fig 3A, but plotted over time. (B) As in (A), but averaged over the eight conditions for each pursuit direction. Error shading = ±1 STE. The curves for the predicted signal are mostly obscured by those for the decoded eye, reflecting the close match. (C) The data from (B) plotted in 2D space (collapsed over time). Time points before the onset of steady pursuit are plotted as smaller circles. (CW = clockwise, CCW = counter-clockwise). The solid black arrow indicates the true eye position, averaged over clockwise and counter-clockwise directions. The dashed curve indicates the performance predicted on the basis of fixation decoding.

An inward bias was expected, however, because a similar pattern of centripetal errors was observed even during the initial *fixation* epoch on these trials, when the eyes were stationary (Fig 3A). Thus, the decoder performance during pursuit might be as we would expect if these same spatial errors manifest as the eye passed through each waypoint. To test this, we generated a *predicted* pursuit curve – under the hypothesis that the neural code for eye position is the same during fixation and pursuit – by interpolating the decoded fixation data points with a 2D cubic spline. Spatially, the decoded eye during pursuit should travel around this circuit (Fig 3A). Over time, however, these spatial biases also translate into *temporal* distortions relative to the actual eye. To understand why, consider that the travel time and distance between one waypoint and the next was approximately constant for the real eye; in contrast, the interpolated distance (along the cubic spline) between one *mis-represented* waypoint and another varied, meaning that the predicted pursuit signal would need to either speed up or slow down to reach the waypoint at the right time. This benchmark curve therefore had idiosyncratic dynamics for each of the sixteen pursuit conditions (Fig 4A, B).

The actual eye position signal decoded from V1 neural activity during pursuit closely matched these fixation-based predictions (*R^2^* = 0.91 and 0.97 for CW and CCW, respectively). This is further evident in Figure 4C, where the decoded positions are plotted over 2D space (collapsed over time) along with those from the predicted pursuit signal. These results show that the representation of eye position in V1 uses a similar code for both stationary and moving eyes.

The updating delay of the decoded signal during pursuit, however, is less clear than it was for saccades. The phase parameter of the sinusoid fit to the averaged pursuit time course suggests that the V1 representation *led* the real eye by 35 ms and 61 ms for CW and CCW pursuit, respectively (consistent with findings in parietal cortex ^35^). However, the decoded trajectories varied considerably across the different starting positions (and pursuit directions), and in many cases were not simply time-shifted versions of the actual eye (i.e. they were not all sinusoidal). These spatial errors can be difficult to distinguish from local shifts in timing, as noted above, complicating the measurement of a simple lag. Indeed, calculating the lag relative to the hypothetical signal predicted from fixation performance (rather than the actual eye), which provided a better match to the decoded pursuit data spatially, gives very different values: a latency of 58 ms for the CW conditions (consistent with the estimate from the saccade task) and −163 ms (i.e. anticipatory) for CCW. The lag during pursuit thus remains unclear.

### How well does V1 code the true positions of visual objects?

We have shown that V1 carries a reliable representation of eye position during exploratory and pursuit movements. What does this mean for the representation of visual space in V1? Geometrically, the true location of an object (o_w_) is the sum of its position on the retina (o_r_) and eye position (e_w_; we ignore head-movements; Fig 1). This means that, by definition, a region of cortex that simultaneously encodes o_r_ and e_w_ implicitly also represents o_w_ (through the availability of o_r_ + e_w_, as long as downstream neurons can extract this information). The presence of a robust place code for o_r_ in V1 is well established: neurons have RFs that remain at a fixed position on the retina irrespective of eye position ^6^. Hence, a point stimulus evokes a narrow hill of activity in the V1 retinotopic map that moves across the cortical sheet according to its own movements and those of the eyes. Our results above show that neurons within a local region of the map also encode e_w_. This implies that V1 as a whole could hold a stable representation of visual space even when the eyes move.

To quantify how well V1 would represent the *true* location of a visual object, we simulated an observer viewing scenarios like those in Figure 1. The eye movements in the simulations were identical to those in our real experiment. The simulated observer located the moving visual object by combining its momentary position on retina, o_r_(t), with the *empirical* eye position we extracted from neural activity in V1 (i.e., Fig 3C and 4B), ê_w_(t), both represented in our analysis as 2D vectors. To take into account the visual latency of our V1 neurons, we lagged o_r_(t) by 53 ms (see Methods). o_r_(t) was thus used as a proxy for the hill of activity this hypothetical object would evoke in the V1 map rather than its position on the retina per se.

This hypothetical observer perceived the movements of objects to be close to their real-world movements, despite grossly distorted trajectories across the retina (and across the V1 map; Fig 5). To benchmark this performance, we compared these reconstructions with a second hypothetical observer who used only the retinal information to locate the object (i.e. by equating retinal coordinates with world-coordinates). We defined a compensation index (CI; see Methods), in which a value of 1 corresponds to a perfect reconstruction of the object’s trajectory and 0 corresponds to the error expected using only its retinal trajectory. Under the traditional view of V1 – in which neurons support only a retinotopic position-code – the CI values should all be close to zero. Our results, however, showed a value of 0.99 for saccades, and 0.90 and 0.93 for clockwise and counter-clockwise pursuit, respectively, showing that this hypothetical observer compensated almost completely for the effects of eye movements.

**Figure 5.**
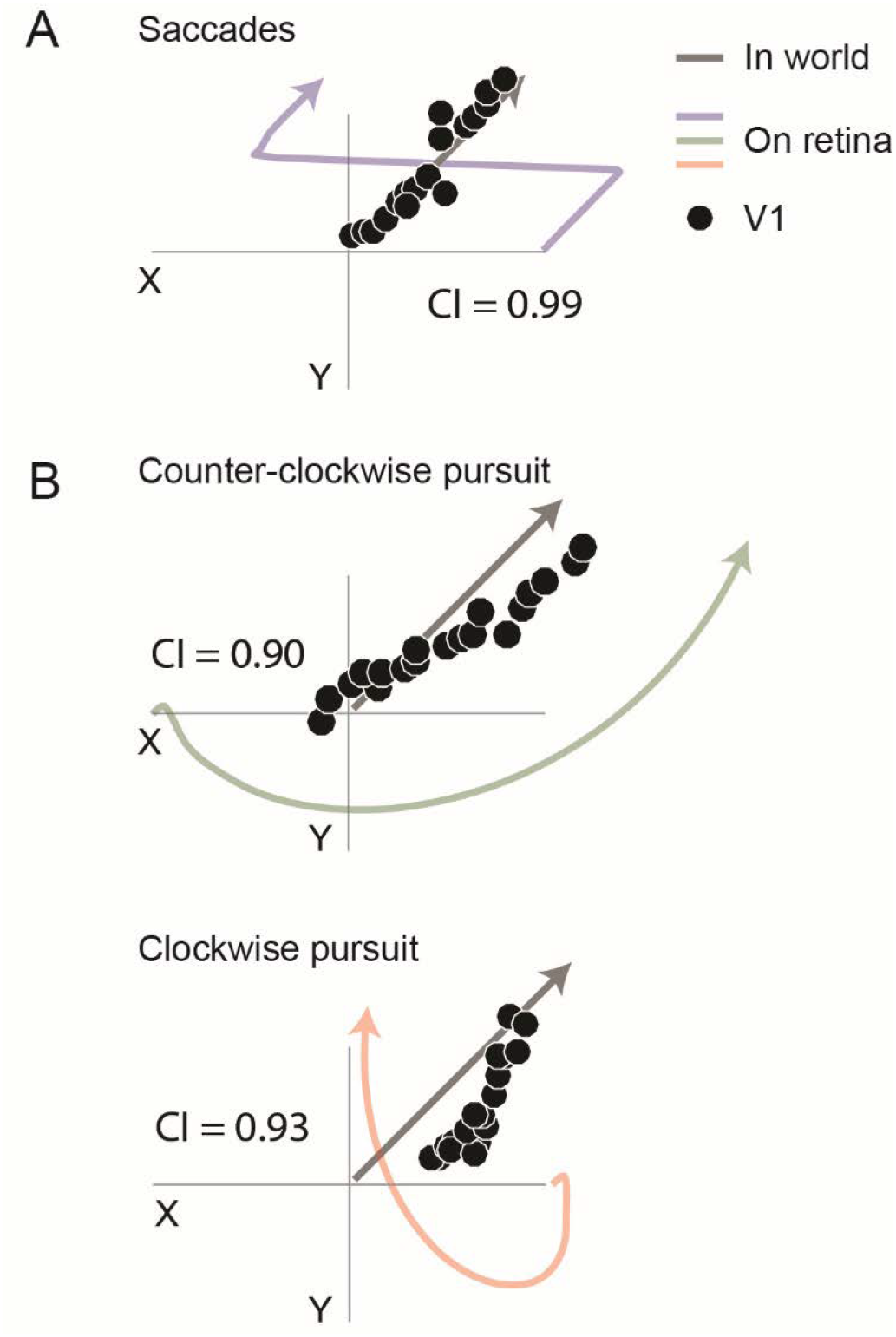
Population codes for eye position in V1 should allow the area as a whole to represent the true locations of objects in the world. We simulated an observer that localises objects by combining their instantaneous positions in the V1 retinotopic map with our experimentally measured eye-position signal. The simulated scenarios were identical to our real experiment – including the saccades and pursuit eye movements – but with the addition of an object in linear motion (black arrow) as in Figure 1. The observer’s “perception” (black circles), which indexes the representation of visual space in V1, was accurate even though its projection on the retina differed greatly under different eye movements (coloured arrows). A compensation index (CI) was calculated (see Methods) in which a value of 1 represents a perfect reconstruction and a value of 0 represents no compensation at all (i.e. using the retinal image alone). (A) A saccade, identical to that in our real experiment, was performed mid-way through the object’s trajectory. (B) Object motion was viewed during clockwise or counter-clockwise pursuit.

## Discussion

Our results show that neurons in primary visual cortex carry a distributed signal that serves as an intrinsic eye tracker, representing the moment-to-moment direction of gaze. This population code was nimble enough to keep up with the eye during exploratory, saccadic eye movements, and also tracked the eye during smooth pursuit movements. Together with V1’s well-established representation of retinal position (i.e. the retinotopic map), this signal provides a plausible neural substrate for the stability of visual perception across eye movements, and suggests that the sensorimotor transformations needed for action begin at the earliest stage of cortical vision.

### Where in the brain is the stable world represented?

Figure 5 showed that V1 contains the information necessary to represent visual space in a way that is much more robust to the effects of eye movements than is suggested by its retinotopic organisation and eye-centred receptive fields ^6^. That is, V1 neurons represent not only the locations of features on the retina (which change every time the eyes move), but also their true locations in the world (technically, their positions relative to the head; Fig 5). By ‘represent’ we mean that an algorithm can extract the information from activity in V1 alone, without accessing any other area. We showed this by construction; the algorithm used in Figure 5 added the 2D vectors of eye position decoded from our data with a (hypothetical, but well-established) 2D vector of a feature’s retinal position.

One might object, however, that neurons do not perform vector addition, and therefore that this algorithm might require a series of operations in downstream areas. If so, it might be better to say that the downstream area represents the stable world, using information retrieved from V1. There is, however, an extensive body of modelling work showing that vector-sum operations can be performed directly through appropriate pooling of activity in gain-field population codes like those seen in V1 ^19, 20, 23, 36^. This operation can be implemented in a single synaptic step and with fixed pooling weights. Hence, our results show that any set of downstream neurons can incorporate the true locations of visual features into their computations.

Moreover, they can do so without regard for the behavioural state of the animal: our decoder was trained using only firing rates observed during fixation, but it was nevertheless able to track the eye across saccades and during pursuit eye movements. This suggests that V1 neurons are modulated by the same source of eye position information during different oculomotor behaviours, and that eye position can be read out with a fixed set of pooling weights. This not only simplifies the read out for vision in real time, but also the challenge of learning such weights during development and maintaining them over time.

Our finding that V1 is home to a representation of visual space does not preclude the existence of similar representations in other visual areas. In fact, we speculate that downstream areas with complex visual form selectivity (V2, V4, IT) represent the world location of those features using similar distributed representations. The presence of eye position signals in these areas supports this view ^2, 37-39^. Similarly, our conclusions are compatible with other purported roles for eye-position information in V1, including the prioritization of visual signals arising from directly in front of an observer’s body ^3, 40^, and the stabilization of image representations during fixational eye movements ^41-44^.

### Robustness in rich visual scenes

In our simulated experiments (Fig 5), and in the classical modelling work ^19, 20, 23, 36, 45^, the visual scenes consisted only of a single, isolated object; in contrast, real visual scenes are cluttered and have high dimensionality (e.g. colour, orientation, luminance, contrast, spatial frequency). This has consequences for the representation of the retinal position, as there is not a single hill in the cortical map but rather a “mountain range” reflecting the structure of the scene. Similarly, it also affects the representation of eye position because the responses of single neurons will be determined by the image features within their receptive fields. This raises the question whether the V1 eye-position signal we have observed is robust in the face of rich visual scenes. Our results, combined with earlier work, suggests that that this should be the case.

First, our experiments used white-noise patterns. These patterns spanned a large set of black-and-white scenes through their random pixel structure, but we were nevertheless able to decode eye position with good accuracy and precision. This was true even though we had access to a vanishingly small number of neurons relative to the total pool in V1. Second, decoding was unaffected by overall changes in the responsivity of the population over the course of individual trials, suggesting a degree of scale invariance (Fig 3C). Finally, we note that multiplexing of eye position information with, for instance, orientation information, is no different from multiplexing two visual feature dimensions. For instance, a single neuron may respond identically to an optimal orientation at a suboptimal spatial frequency (or colour, contrast, etc.) or a suboptimal orientation at an optimal spatial frequency. In other words, even the representation of orientation in individual V1 neurons is highly ambiguous and can only be resolved at the population level. Our results show that eye position information should be viewed similarly, as a feature of the visual input that is represented across a population of neurons. The fact that changes in eye-position typically alter only the *gain* of neural responses in V1, and not their selectivity to visual features, should facilitate the separation of eye-position signals ^1, 46^. Alternatively, they could be separated by making use of the ^~^50% of neurons that do not have gain-fields (as observed here, and in ref ^1^), possibly by a pairwise subtraction of neural pools with comparable visual selectivities.

### Beyond V1

Computations that combine visual and eye-position signals are obviously simplified when such signals are aligned temporally. This is particularly important in the case of saccadic eye movements, where the eye moves with very high velocity across large distances within a scene. Our simulations (Fig 5) shows that temporal alignment was sufficient to compute the true positions of objects in the world. Indeed, our best estimate of the latency of the eye position signal (45 ms) – from the saccade task – closely matched the visual latency in these neurons (53.11 ms; SD = 9.93 ms; see Methods). Faster still was the representation of the eye during pursuit eye movements: our estimates hinted at a predictive signal, consistent with findings in parietal cortex ^27, 28, 35^, though in the pursuit case our estimates were inconsistent across conditions and likely corrupted by imperfect spatial coding of the pursuit trajectory. Regardless, the V1 eye position signal appears to be updated earlier than the proprioceptive representation of the eye in primary somatosensory cortex ^48, 49^. This suggests that it arrives via a more direct pathway, potentially even embedded within the incoming visual information from the lateral geniculate nucleus ^33, 50-52^, consistent with earlier findings in V1 ^1^.

We speculate that the cortical eye position signal arrives first in V1 and then propagates through the cortical hierarchy along with the feedforward visual information. This would lead to progressively later lags as the combined signals reach later stages of the cortical pathway, consistent with measurements in parietal cortex ^53^, while naturally maintaining the spatiotemporal alignment needed for stability within each area (and hence at every spatial scale). In parietal cortex, these signals likely interact with other mechanisms that contribute to stability, including representations of eye position and retinal position (e.g. so-called “remapping”) that are updated even *before* the eyes move ^14, 17, 18, 27, 28^.

In conclusion, we suggest that the accurate and nimble representation of eye position at this first stage of cortical vision – which feeds both the dorsal and ventral processing streams – is a critical ingredient in the computations underlying spatial stability. Our simulations show that this signal alone could explain why objects do not appear to change positions every time we move our eyes. Further, it could play a similar role in the continuity of action planning, object recognition, and decision-making processes across eye movements.

## Online Methods

### Subjects

Behavioural and electrophysiological measurements were performed in two adult, male macaques (*Macaca Mulatta*; M1: weight = 14 Kg; M2: weight = 9.9 Kg). All procedures conformed to the National Institute of Health guidelines for the humane care and use of laboratory animals and were approved by the Rutgers University Animal Care and Use Committee.

### Animal preparation

Surgery was performed under aseptic conditions and general anaesthesia (isoflurane), and analgesics (ibuprofen and buprenorphine) and antibiotics were delivered during recovery. Animals were first fitted with a titanium head-post (Gray Matter Research) to stabilize the head during recording sessions using standard surgical procedures. After recovery, and completion of simple fixation training, animals were prepared for chronic, subdural implantation of one (M2; left-hemisphere) or two (M1; right-hemisphere) 32-channel, planar microelectrode arrays (Floating Microelectrode Arrays; MicroProbes for Life Science) into the parafoveal region of V1. Each array consisted of a 4 × 8 arrangement of platinum/iridium electrodes (0.6–1.5 mm in length [tiered]), spanning 1.2 × 3.4 mm and separated by a distance of 0.44 mm (0.8–1 MΩ impedance at 1 kHz). A craniotomy was performed immediately anterior to the occipital ridge and lateral from the midline, and a small patch of dura was removed to provide access to the parafoveal region of V1. Each array was implanted normal to and flush with the cortical surface using manual descent and depth control via a stereotax-mounted micromanipulator ^54^. Neural signals were communicated via a fine bundle of wires that was routed out and along the margin of the craniotomy to Omnetics connector(s) mounted in a stainless steel chassis (M2) or dental acrylic (M1). The two arrays in M1 were implanted directly side-by-side in the same hemisphere. Full closure of the craniotomy was achieved using a layer of DuraGen (Integra) and dental acrylic to reduce risk of infection (no subsequent signs of infection were observed). The animals were allowed to fully recover before the commencement of electrophysiological recordings.

### Stimuli and apparatus

Visual stimuli were presented on a CRT display (Sony GDM-520, NVidia 8800 GT graphics card) at a vertical refresh rate of 150 Hz and resolution of 1024 × 768. The visible display area subtended 43.7° × 33.6° at a centre-aligned viewing distance of 50 cm and was linearised for luminance at [CIEx, CIEy] = [0.33, 0.33]. Stimulus presentation and experimental flow was controlled by custom software written in C++ (Neurostim, available at https://sourceforge.net/projects/neurostim/; last accessed January, 2018). The animal was seated in a primate chair with its head stabilized using a head-post. During experimental trials, neurons were stimulated visually by dynamic binary noise, in which every pixel in the display was switched to black (0.6 cd/m^2^) or white (50 cd/m^2^) randomly and independently on every second retrace of the display (and maintained its value for the intervening frame). The mean background luminance was therefore 25.3 cd/m^2^. Fixation and pursuit targets were red spots (diameter = 0.25°) placed atop a larger black spot (diameter = 0.63°) that masked the surrounding noise pixels. Left and/or right eye position was recorded using an infrared video-based eye tracker (Eyelink II, SR Research) at a sample rate of 250 or 500 Hz, interpolated offline to 1 kHz, and temporally aligned to the neural data.

### Behavioural tasks

Each trial began with a fixation target presented at one of nine positions (eight equally spaced around a virtual circle [radius = 7.5°] and one in the centre of the display) against a grey background (25.3 cd/m^2^; Fig 2A). When ready, the animal fixated the target to initiate the trial proper. The onset of fixation triggered the presentation of binary noise across every pixel in the display, which continued until the end of the trial. On saccade trials, the animal maintained fixation for 1100 ms before the target stepped to the diametrically opposite position on the circle. The animal was required to saccade to the target within 450 ms and hold fixation for a further 1100 ms. On pursuit trials, the target started to move after animal had maintained initial fixation for 500 ms, and then moved at constant speed (10°/s [i.e. degrees of visual angle]) around the circle (clockwise or counter-clockwise) to the diametrically opposite position (at which time the trial ended). In the case of the central fixation target, the animal was required to maintain fixation for the duration of the trial. Trials in which the animal failed to maintain gaze within 1.5° of the target were terminated immediately without reward, while successfully completed trials were rewarded with diluted fruit juice. The screen returned to grey during the 1200 ms inter-trial interval and the animal was free to look in any direction. Animals M1 and M2 completed an average of 71.26 (SD=16.88) and 50.90 (SD=23.69) saccade trials correctly per starting position, per session. They also completed an average of 36.51 (SD=7.95) and 21.46 (SD=4.87) pursuit trials correctly per starting position, per pursuit direction (clockwise, counter-clockwise), per session.

### Electrophysiology and data processing

Signals from the V1 microelectrode array were digitized (14 bit at 25 kHz) and band-passed (Butterworth, 120 Hz at 12 dB/octave to 6 kHz at 24 dB/octave) using AlphaLab recording hardware (Alpha Omega Co.). Candidate action potentials were detected using a threshold of 3.5 standard deviations separately for each electrode and sorted into clusters using an automated algorithm (superparamagnetic clustering ^55^). Each cluster was categorized by hand as a single-unit, multi-unit, or junk on the basis of its (1) mean cluster waveform; (2) inter-spike interval histogram, and (3) visual responsivity (assessed by peristimulus time histograms of spike counts aligned to the onset of the binary noise). All analyses were performed on single- and multi-units without distinction unless otherwise specified.

### Estimation of receptive fields

The linear receptive-field (RF) of each unit was estimated from the fixation/saccade trials using a standard spike-triggered average (STA). The approximate locations of the RFs were known from preliminary analyses and previous recordings. Accordingly, the STA was calculated only for noise pixels within a 10.7° × 10.7° (251 x 251 pixels) square region of interest (ROI) centred on the position [−4°, −1°] relative to the fixation point (i.e. from different screen locations depending on the fixation location). Each pixel subtended 2′ 33″ (0.04°) of visual angle.

The luminance values (coded as 0 [black] and 1 [white]) of every pixel were collated into an *m* × *n* matrix, *L,* where *m* is the number of pixels in the ROI and *n* is the total number of frames (pooled across time and all trials). The spike times for each unit were aligned to the first frame of the stimulus and binned by counting the number of spikes that occurred within each stimulus frame (i.e. 13.33 ms bins). To take into account visual latency, the resulting *n*-length column vector, *C*, was replicated 11 times, each with an additional frame of time lag (i.e. shifted from 13 to 147 ms), and compiled into an *m* x 11 matrix. The space-time STA (*m* x 11) was then calculated through matrix multiplication (*L*C)*. The strongest STAs were found in the three lags from 54 to 80 ms prior to the time of the spike, hence we calculated the spatial receptive field maps in Figure 2 by averaging these three time lags and smoothing with a 2D Gaussian filter (5 x 5 pixels, SD = 2). These visual latencies estimated using the STA approach were consistent with those estimated from onset responses to the noise patterns (53.11ms) using the method of Friedman and Priebe (1998); that is, using only data from 150 ms before to 200 ms after stimulus onset ^56^.

To identify pixels in the STA that were statistically significant, we created an empirical null distribution of STA magnitudes (including the effects of smoothing and other averaging) by repeating our analysis across the entire 1024 x 768 stimulus. These 786,432 pixels could be used as a measure of sampling variability because we knew from previous recordings that almost of them were well outside the tiny V1 RFs. Pixels in the ROI STA were deemed significant if their absolute value exceeded 3.5 times the standard deviation of this null distribution (i.e. *p<0.0002)*. The fact that a small subset of pixels was inside the RF made this a conservative estimate.

To summarize the RFs across all units in an animal (Fig 2C), we calculated population RF densities by convolving each unit’s significance map (where significant and non-significant pixels had values of one and zero, respectively) with a 2D Gaussian filter (10 x 10 pixels, SD = 4) and then summing across all units. Pixels in individual STAs that were false positives would occur at random positions and thus would tend not to accumulate across units and be largely absent in the population density image.

### Eye position gain-fields

Gain-fields were estimated from the fixation/saccade trials. Spike times were aligned to the offset of the primary saccade from the first fixation position to the opposite side of the circle. This eye movement was found offline and defined as the first saccade (detected by the Eyelink system) after the change in target position that had an amplitude of at least 75% of the required distance (15°). Spike times on central fixation trials, which did not require a saccade, were aligned to the mean saccadic offset time from the saccade trials (i.e. the time of the target step plus mean saccade latency and duration) to maintain consistent alignment. Spike times were then binned into 100 ms windows stepped in 25 ms increments to provide a spike-count time course for each trial.

Gain-fields are defined as a systematic change in the mean firing rate of a visual neuron as a function of fixation position despite constant stimulation of the receptive field. Accordingly, we calculated the mean spike-count during the initial fixation period (from −700 ms to −400 ms, equivalent to −570 to 870 ms after the onset of the visual noise on average) separately for each of the nine fixation positions. A statistically significant gain-field was identified using stepwise regression, in which a second–order polynomial was used to model the effect of eye-position (azimuth [X] and elevation [Y]) on mean spike counts, *ĉ*(Equation 1). Coefficient estimates and the associated F-tests were performed using the “stepwiseglm” function in Matlab (Mathworks Inc.). To prevent cardinal biases all terms were retained for decoding (see below), even if the coefficients were individually non-significant ^24^.

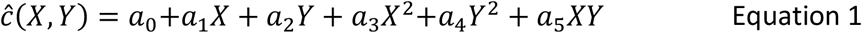

### Population decoding of eye position

Gain-fields provide an intuitive quantitative link between the activity of a neuron and eye-position. Such a link can form the basis of decoding algorithms that predict or estimate eye position using only neural activity across one or more neurons. We have used this approach successfully to decode eye position from populations of neurons across many regions in the dorsal visual system during stationary gaze ^24^, and to examine the dynamics of these representations when the eyes move ^27, 28^ (see also ^35^). In one such study ^24^, we developed a decoding approach that estimates the *most likely* eye position given a set of observed spike counts across a sample of neurons. The method is built on the well-established statistical approach of maximum-likelihood estimation, which takes into account not only how the mean firing rates depends on eye position, but also the associated variability (sometimes referred to as neural “noise”). Here we apply this decoding approach to all V1 units that had significant gain-fields (219/397). The full method has been described in our previous work ^24^ but is summarized here.

### Maximum-likelihood estimation

The traditional definition of a gain-field quantifies only how the mean firing rate of a neuron depends on eye position and says nothing about spike-count variability. We introduced a richer statistical quantification known as a *probabilistic gain-field* (pGF)^24^. Given an eye position, the relative likelihood of observing each possible spike-count (i.e., *c* = 0, 1, 2, 3, etc.) is captured by a probability density function (pdf), *P*(*C*|*x,y*). Our approach is to estimate this generative pdf at each of the experimentally measured eye positions and then to interpolate (in 2D) to unmeasured fixation positions. By doing so with a continuous expression, we can express *P*(*C*|*x,y*) at arbitrary eye positions within or near the sampled range – this representation is the pGF. For decoding, however, we want to come from the other direction: given an observed spike-count, we want to know the relative probability of all possible eye positions, *P*(*X*,*Y*|*c*). This *a posteriori* density can be computed from the pGF using Bayes Rule (we assumed a uniform prior over all eye positions) and then used to generate a point estimate by choosing the most likely current eye position.

The approach extends naturally to a population of neurons. With a pGF in hand for each of *N* recorded V1 units, a vector of observed spike-counts across the sample, *c_pop_*, can be converted into a population estimate of *P*(*X*,*Y*|*c_pop_*) by combining the information across units, implemented as the sum of the (log) posterior probabilities:

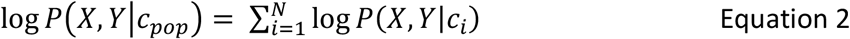

The final estimate of eye-position is taken as the position associated with the highest likelihood, [*x_max_*, *y_max_*].

For this study, we decoded across a discrete grid of possible eye positions spanning ±30° in 0.5° steps both horizontally (azimuth) and vertically (elevation). The spike-count vector, *c_pop_*, was compiled by taking a random trial for each unit from a common experimental condition and treating them as if they occurred simultaneously. This pseudo-population response approach was preferred because our data set included only 7.49 visually-responsive units per recording session on average (SD = 5.35), which was insufficient to decode the eye reliably for all positions. A consequence of this randomization, however, is that it removes correlated variability across the “population” of neurons, which can influence the information that is available to a decoder (for better or worse, depending on its structure) ^57, 58^. Our estimates of decoder precision (i.e. moment-to-moment variability) thus incorporate this potentially incorrect assumption of independence and should be interpreted accordingly.

As described above, we needed an estimate of *P*(*C*|*x*,*y*) at each measured eye position. Ideally, we would use the empirically observed distribution of spike counts for a given unit for this purpose. In studies of this kind, however, it is usually not possible to perform enough measurements to estimate the underlying probability model reliably, particularly for rare spike-count values (which can then cause instability in the decoder). Instead, as in previous work^24, 25^ we made the simplifying assumption that the variability can be approximated by Poisson distributions, in which the mean and variance are equal. This has the convenient property that the traditional gain-field function, which quantifies how mean evoked activity varies with eye position, can be reinterpreted as a continuous expression of the value of the single-parameter (*λ*) Poisson pdf. Accordingly, Equation 1 was used as a generative model for the pGF of each unit.

### Decoding during fixation

The previous section described how we built the decoder from the gain-fields of each recorded unit. To assess how well V1 represented the eye during fixation, the decoder was applied to spike-counts in 100 ms wide windows during the same interval that was used to fit the gain-fields (i.e., −700 ms to −400 ms relative to saccade offset). This was achieved using leave-one out cross-validation, in which all but one trial from each of the nine eye-positions was used to fit the gain-fields (and hence, produce a pGF for each unit). The resultant decoder was then applied to the remaining trial from each condition and the estimated eye position was recorded. This was repeated 1000 times to produce a distribution of estimates for each eye position. The 2D median of each of these distributions (plotted in Fig 3A) was used to summarize the accuracy of the V1 representation, and the span of the middle 50% of the data (horizontally and vertically) was used to represent its precision (error bars in Fig 3A).

For all subsequent analyses (described below) we used a decoder built from this initial fixation interval but without leaving out any trials. This “fixation” decoder was then locked down – no further “learning” could take place – and applied to either different time points to which it was naïve (e.g. during the saccade), or to different trial types (e.g. pursuit trials).

### Decoding across saccades

One-thousand pseudo-population responses were compiled for each starting position in the saccade task and then decoded using the decoder described above. Each time point, from 900 ms before the saccade to 900 ms after, was decoded sequentially and independently and with no knowledge of the true eye position. Hence, only changes in the activity of V1 neurons over time could lead to changes in the decoded eye positions. The result was a distribution of 1000 decoded time courses for each condition. As for the fixation analysis, we used the median and width of these distributions at each time point to assess the accuracy and precision of the decoder. To summarize overall performance, we combined all conditions by rotating them onto a common starting position (the 9 o’clock position) and then taking the mean (of the medians; Fig 3C).

Of key interest was the timing and speed with which the recorded units transitioned from a representation of the initial fixation to that of the final position at the diametrically opposite position on the circle. Like the real eye, the decoded eye showed a sigmoid-like change over time in the saccading channel, while remaining relatively stationary in the orthogonal channel. To quantify this transition, we fitted a cumulative Gaussian (using Matlab’s “lsqcurvefit” function) to the saccade channel. The mean parameter (*μ*) of the fit corresponded to the mid-point of the represented saccade and so was used as the time at which updating occurred. The duration was taken as the interval between the 5^th^ and 95^th^ percentile (i.e. saccade duration = 2*1.65\**σ*). The same method was used to quantify the real eye, which was also binned in 100 ms for consistency with the neural data. This binning dampens the signals, which is why the average saccade duration reported here (114 ms) is longer than its true duration (33.1 ms). This does not affect our conclusions because identical binning was used for the eye and neural data. The difference between the two updating times (*σ_decoded_* – *σ_real_*) was used as the estimated latency of the eye-position signals represented in V1 (i.e. the delay from a true change in eye position to when that change is represented in cortex).

### Decoding during smooth pursuit

The spike times on pursuit trials were aligned to the onset of the target motion and binned in the same way as for saccade trials. Because pursuit trials were not included in all recording sessions, some units had to be removed from the decoder. In all other respects, however, the decoder remained unchanged and hence had never been exposed to a single data point from pursuit trials. We decoded the neural data in the same manner as described above for saccade trials and extracted the same measures of accuracy and precision over time.

Both the real and decoded eye position traced out approximate sine/cosine functions over time for the horizontal and vertical channels. We fitted such functions (Equations 3 and 4) to the averaged decoded time courses (Fig 4B) and to the real eye and used the difference in phase between them to estimate the latency of the V1 eye-position signal. Data from the first 400 ms after the target motion onset (where pursuit was not yet stable) was excluded.

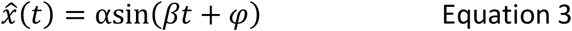

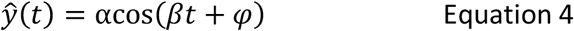

To ensure that all temporal differences manifest in the phase (*φ*), the period parameter (*β*) was constrained to be the same for the real and decoded eye position traces. This was achieved by first fitting Equations 3 and 4 to the two data sets separately, and then re-fitting them with *β* fixed to the average of the two values from the first stage.

Decoder performance during the initial fixation period of pursuit trials (i.e. –200 ms to +100 ms relative to target motion onset) was examined for multiple purposes. First, it allowed us to compare the spatial errors with those of the saccade trials for this reduced population size (Fig 3A). Second, and more importantly, we used these errors to construct baseline time courses for how the decoder should perform during smooth pursuit *if the neural code for eye position remained identical to that of a stationary eye* – that is, if neural firing rates depended only on the instantaneous eye position (Fig 4).

Each pursuit condition passed through five of the positions measured during fixation (including the start- and end-points) during its half-circle trajectory, which we refer to as “waypoints”. To generate expected trajectories for each condition, we first fitted a 2D cubic spline through the eight decoded positions associated with each waypoint (i.e. the dotted line in Figure 3A). Spatially, the predicted pursuit signal should travel around this spline, but how it should do so in time is complex and requires explication.

As a hypothetical example, suppose the decoded eye position for the 9 o’clock position during fixation was 1° above the true position. What would happen if this same error occurred during pursuit at the moment the eye passed *through* the 9 o’clock waypoint? The decoder would appear to *lead* the eye during clockwise pursuit because the decoded position would already be ^~^1° ahead of the upward moving eye. Conversely, the same error would look like a temporal *lag* when the eyes were moving counter clockwise because the decoded position would remain ^~^1° behind the downward moving eye. Thus, although the real eye travels from one waypoint to the next with approximately constant speed and distance, the signal that is predicted on the basis of the fixation performance should travel different distances depending on the errors associated with the departure and arrival waypoints.

With this in mind, we computed a predicted pursuit signal by (1) noting the times at which the eye crossed each waypoint, and (2) moving around the cubic spline from the (erroneous) position associated with each waypoint to that of the next within the same time interval as the real eye. These predictions are plotted in Figure 4.

### Assessing visual stability

A neural code for vision is stable if it remains spatially accurate even when the eyes move. In the current context, this requires encoding an object’s position relative to the head (or equivalently, because the animal was head-fixed, relative to the display), because this position remains the same irrespective of eye position. To a good approximation, the head-centred location of an object is equal to the sum of two quantities: its projected position onto the retina (i.e., its horizontal and vertical distance from the fovea, in degrees of visual angle), and eye position (i.e. the horizontal and vertical angles of rotation in the orbit). Assuming a stationary object, both of these quantities change every time the eyes move but their sum remains constant.

Given that objects move in the real world, however, all terms in this linear relation are functions of time. Moreover, the cortical representation of an object’s position on the retina, *r̂*, presumably encoded via retina-fixed receptive fields ^6^, does not reflect its current position, but rather where it was some time ago, reflecting the transmission delay (visual latency, *V_L_*) from the retina to the cortex. That is, assuming a perfect cortical code:

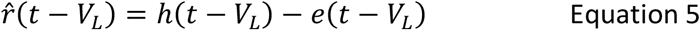

where *h* is the true location of the object in the world and *e* is the true eye position.

Similarly, the brain does not have access to eye position directly but must estimate it from an internal signal (e.g. efference copy from the motor system or proprioception originating in the eye musculature). To maintain spatiotemporal alignment with the incoming visual information, this internal eye-position signal, *ê*, must but also lag behind the eye by the visual latency (as well as faithfully representing the dynamics of the eye). In that case, the combination of retinal and eye-position signals gives rise to a stable internal representation of the outside world (at least, as it was *V_L_* milliseconds ago).

To assess visual stability in V1 within this framework, we simulated an object moving across the display during the saccade and pursuit tasks, and used the observed V1 data to reconstruct a neural estimate of its position over time. The simulated object moved at a constant velocity in the fronto-parallel plane, which varied in speed and direction across simulated trials (8 evenly spaced directions between 0 and 2π, 5 speeds [0°/s, 2°/s, 4°/s, 8°/s, 16°/s]). A neural estimate of the object’s position was calculated by combining the decoded eye position signal (i.e. the time course shown in Figure 3C or Figure 4B) with an idealised representation of the object’s position on the retina. The latter was calculated as the difference between the position of the simulated object and the true eye position (as reported by the eye tracker, also shown in Figure 3C). This representation of the retinal projection was then lagged by the visual latency.

The two quantities – visual and eye-position – were represented as [x,y] vectors and summed to provide an estimate of the object’s position at each moment in time, across saccades and during pursuit. That is:

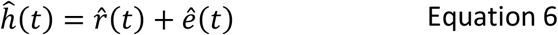

We compared the estimated and true object positions to quantify visual stability in V1. We first calculated a traditional sum-of-squared errors to quantify the spatial mismatch (i.e., 2D Euclidean distance) between the predicted object trajectory and its true trajectory:

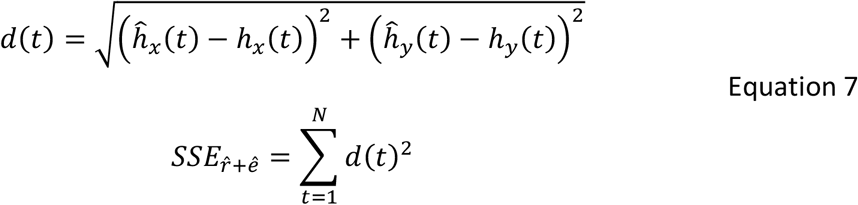

To provide a meaningful reference, we compared this error value with that observed if we used only the retinal signal for localisation (i.e. only the object’s position within the V1 map) and ignored the eye-position signal. That is:

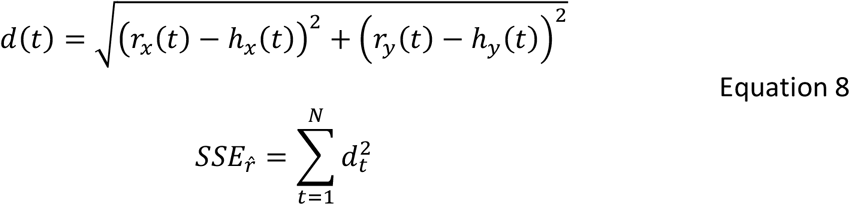

We defined a compensation index (CI) to capture how well gain-fields allowed V1 to compensate for the effects of eye movements:

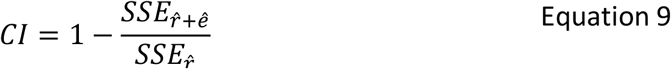

The index has a value of zero if the prediction is no better than using V1 receptive fields alone, and 1 in the case of perfect prediction (the upper-bound). The index has no lower-bound (negative values are possible).

The CI index was computed for all of the simulated object trajectories during saccades, and during both pursuit directions (clockwise and counter-clockwise).

## Acknowledgements

This work was supported by the National Health and Medical Research Council of Australia (A.P.M.; APP5525487 and APP1083898) and the National Institutes of Health, USA (B.K.; EY017605 and MH111766). The contents of the published material are solely the responsibility of the Administering Institution, a Participating Institution, or individual authors and do not reflect the views of the NHMRC or the NIH. The authors acknowledge Anne McCormick for veterinary and technical assistance, Till Hartmann for help with the experimental set-up, and Maria del Mar Quiroga, Jacob Duijnhouwer, and Shaun Cloherty for useful discussions.

## Author contributions

A.P.M and B.K. performed all aspects of this work, including conceptual development and experimental design, surgical procedures, animal training, programming for experimental control and analysis, and preparation of the manuscript.

